# Models of heterogeneous dopamine signaling in an insect learning and memory center

**DOI:** 10.1101/737064

**Authors:** Linnie Jiang, Ashok Litwin-Kumar

## Abstract

The *Drosophila* mushroom body exhibits dopamine dependent synaptic plasticity that underlies the acquisition of associative memories. Recordings of dopamine neurons in this system have identified signals related to external reinforcement such as reward and punishment. However, other factors including locomotion, novelty, reward expectation, and internal state have also recently been shown to modulate dopamine neurons. This heterogeneity is at odds with typical modeling approaches in which these neurons are assumed to encode a global, scalar error signal. How is dopamine dependent plasticity coordinated in the presence of such heterogeneity? We develop a modeling approach that infers a pattern of dopamine activity sufficient to solve defined behavioral tasks, given architectural constraints informed by knowledge of mushroom body circuitry. Model dopamine neurons exhibit diverse tuning to task parameters while nonetheless producing coherent learned behaviors. Our results provide a mechanistic framework that accounts for the heterogeneity of dopamine activity during learning and behavior.

## Introduction

Dopamine release modulates synaptic plasticity and learning across vertebrate and invertebrate species^1,2^. A standard view of dopamine activity, proposed on the basis of recordings in the mammalian midbrain dopaminergic system, holds that dopamine neuron firing represents a “reward prediction error,” the difference between received and predicted reward^3^. This view is consistent with models of classical conditioning experiments and with reinforcement learning algorithms that learn to choose the most rewarding sequence of actions^4^. A frequent assumption in these models is that the scalar reward prediction signal is globally broadcast to and gates the modification of synaptic connections involved in learning. However, studies in both vertebrates and invertebrates suggest that dopamine neuron activity is modulated by other variables in addition to reward prediction error, and that this modulation is heterogeneous across populations of dopamine neurons^5^.

Early studies in arthropods identified roles for dopamine in a variety of functions^6–11^. In *Drosophila,* both memory^12^ and other functions including locomotion, arousal, sleep, and mating have been associated with dopamine signaling^11^. Associative olfactory learning in *Drosophila* requires a central brain area known as the mushroom body^13–15^, and many studies of dopamine neurons innervating this area have focused on activity related to reward and punishment and its roles in the formation of appetitive and aversive memories^16–22^. In the mushroom body, Kenyon cells (KCs; green neurons in Fig. 1A) conveying sensory information, predominantly odor-related signals, send parallel fibers that contact the dendrites of mushroom body output neurons (MBONs; black neurons in Fig. 1A). The activation of specific output neurons biases the organism toward particular actions^23,24^. Output neuron dendrites define discrete anatomical regions, known as “compartments,” each of which is innervated by distinct classes of dopaminergic neurons (DANs; magenta neurons in Fig. 1A). If the Kenyon cells and dopamine neurons that project to a given output neuron are both active within a particular time window, KC-to-MBON synapses are strengthened or weakened depending on the relative timing of Kenyon cell and dopamine neuron activation^25–28^. The resulting synaptic modifications permit flies to learn and update associations between stimuli and reinforcement.

**Figure 1:**
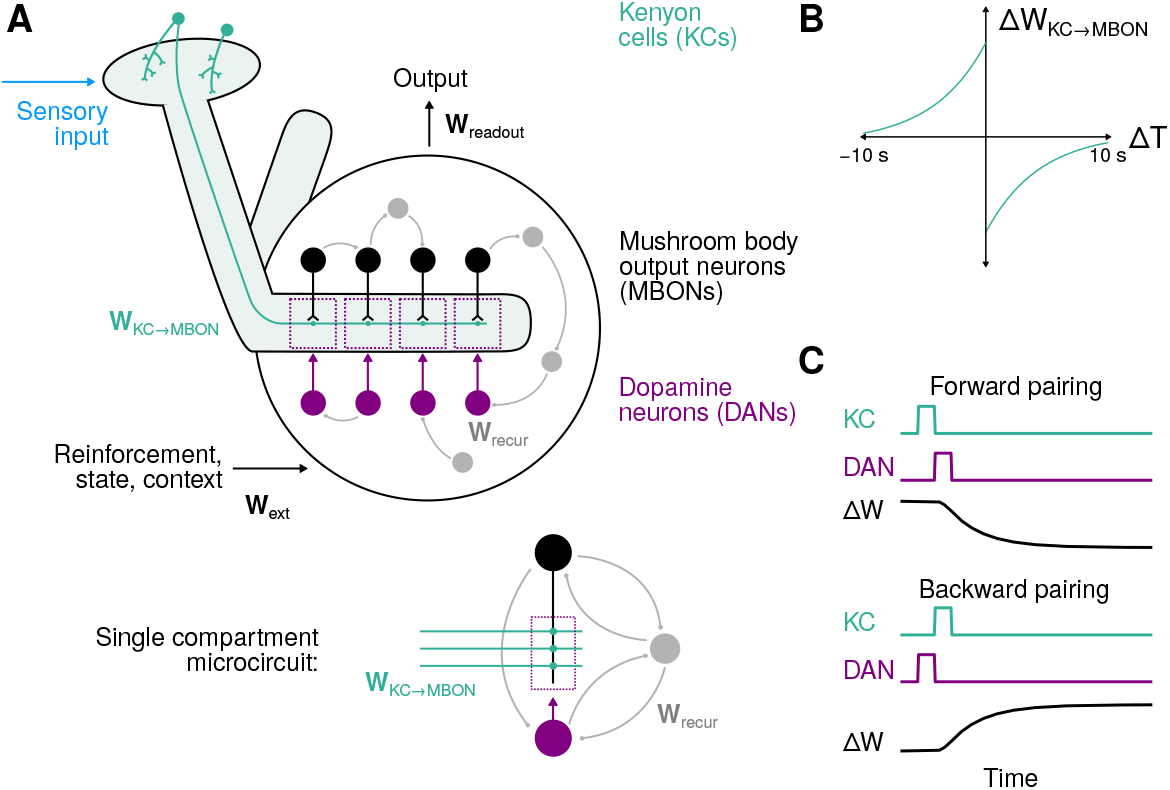
Diagram of the mushroom body model. **(A)** Kenyon cells (KCs) respond to stimuli and project to mushroom body output neurons (MBONs) via weights **W**_KC→MBON_. These connections are dynamic variables that are modified according to a synaptic plasticity rule gated by dopamine neurons (DANs). Output neurons and dopamine neurons are organized into compartments (dotted rectangles). External signals convey, e.g., reward, punishment, or context to the mushroom body output circuitry according toweights **W**_ext_. A linear readout with weights **W**_readout_ is used to determine the behavioral output of the system. Connections among output neurons, dopamine neurons, and feedback neurons (gray) are determined by weights **W**_recur_. Inset: expanded diagram of connections in a single compartment. **(B)** The form of the dopamine neuron-gated synaptic plasticity rule operative at KC-to-MBON synapses. Δ*T* is the time difference between Kenyon cell activation and dopamine neuron activation. **(C)** Illustration of the change in KC-to-MBON synaptic weight Δ*W* following forward and backward pairings of Kenyon cell and dopamine neuron activity.

In addition to classical reward and punishment signals, recent studies have shown that variables including novelty^29^, reward prediction^30–32^, and locomotion-related signals^33^ are encoded by mushroom body dopamine neurons. In mammals, dopamine signals related to movement, novelty and salience, and separate pathways for rewards and punishment have also been identified in midbrain regions^5,34–42^. These observations call for extensions of classic models that assume dopamine neurons in associative learning centers are globally tuned to reward prediction error^43^. How can dopamine signals gate appropriate synaptic plasticity and learning if their responses are modulated by mixed sources of information?

To address this question, we develop a modeling approach in which networks that produce dopamine signals suited to learning a particular set of behavioral tasks are constructed. Our key methodological advance is to augment standard recurrent neural network models, which employ fixed synaptic weights to solve tasks after optimization ^44^, with synapses that exhibit fast dopamine-gated plasticity via an experimentally determined plasticity rule^28^. We employ a “meta-learning” approach involving two phases^45^ (Supplemental Fig. 1). First, we optimize the network connections responsible for producing suitable learning signals in dopamine neurons. Next, after these connections are fixed, we examine the network’s behavior on novel tasks in which learning occurs only via biologically plausible dopamine-gated plasticity. Due to the well-characterized anatomy of the mushroom body and knowledge of this plasticity rule, our approach allows us to generate predictions about the activity of multiple neuron types^28,46^. Comprehensive synapse-level wiring diagrams for the output circuitry of the mushroom body will soon be available, which will allow the connectivity of models constructed with our approach to be further constrained by data^47–50^. As the dynamics of our models, including the dopamine-gated plasticity, are optimized end-to-end only for overall task performance, our model predictions do not require a priori assumptions on what signals the dopamine neurons encode. In particular, our methods do not assume that each dopamine neuron carries a reward prediction error and instead allow for heterogeneous activity across the dopamine neuron population.

The meta-learned networks we construct are capable of solving complex behavioral tasks and generalizing to novel stimuli using only experimentally constrained plasticity rules, as opposed to networks that require gradient descent updates to network parameters to generalize to new tasks. They can form associations based on limited numbers of stimulus/reinforcement pairings and are capable of continual learning, which are often challenging for artificial neural networks^45,51^. In the models, different dopamine neurons exhibit diverse tuning to task-related variables, while reward prediction error emerges as a mode of activity across the population of dopamine neurons. Our approach uncovers the mechanisms behind the observed heterogeneity of dopamine signals in the mushroom body and suggests that the “error” signals that support associative learning may be more distributed than is often assumed.

## Results

### Modeling recurrent mushroom body output circuitry

The diversity of dopamine neuron activity challenges models of mushroom body learning that assume dopamine neurons convey global reward or punishment signals. Part of this discrepancy is likely due to the intricate connectivity among output neurons, dopamine neurons, and other neurons that form synapses with them^46,50^. We therefore modeled these neurons and their connections, which we refer to collectively as the mushroom body “output circuitry,” as a recurrent neural network (Fig. 1A). This model network consists of 20 output neurons, 20 dopamine neurons, and 60 additional recurrent feedback neurons. Recurrent connections within the network are defined by a matrix of synaptic weights **W**_recur_. Connections between all of these 100 neurons are permitted, except that we assume connections from dopamine neurons to output neurons are modulatory and follow a compartmentalized organization (see below; Fig. 1A, inset). Synapses from 200 Kenyon cells onto output neurons provide the network with sensory information and are represented by **W**_KC→MBON_. Kenyon cells do not target neuron types other than output neurons in our model. Separate pathways convey signals such as reward or punishment from other brain regions, via weights **W**_ext_. The dynamics of the *i*th neuron in our model of the output circuitry are given by:

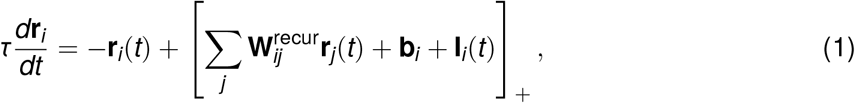

where [·]_+_ represents (elementwise) positive rectification. The bias **b**_*i*_ determines the excitability of neuron *i*, while **I**_*i*_(*t*) represents its input from non-recurrent connections (Kenyon cell input via **W**_KC→MBON_ in the case of output neurons, and external input via **W**_ext_ in the case of feedback neurons; see Methods). We do not constrain 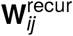, except that entries corresponding to connections from dopamine neurons to output neurons are set to zero, based on the assumption that these connections modulate plasticity of KC-to-MBON synapses rather than output neuron firing directly (see Discussion).

The objective of the network is to generate a desired pattern of activity in a readout that represents the behavioral bias produced by the mushroom body. The readout decodes this desired output through a matrix of weights **W**_readout_. In our first set of experiments, this readout will represent the one-dimensional valence (appetitive vs. aversive) of a stimulus decoded from the output neurons (meaning that **W**_readout_ is a 1 × *N*_MBON_ matrix; later, we will consider more sophisticated readouts):

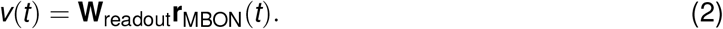

To achieve the task goal, trials are randomly generated and the following objective function, which depends on the parameters of the network **θ** and represents the loss corresponding to an individual trial consisting of *T*timesteps, is minimized through stochastic gradient descent:

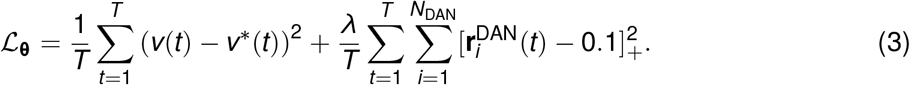

The first term represents the difference between the decoded valence and a target valence *v*\* that is determined by the task being learned. The second term is a regularization term that penalizes dopamine neuron activity that exceeds a baseline level of 0.1 (in normalized units of firing rate and with *λ* = 0.1). We verified with simulations that the regularization term does not significantly affect overall network performance.

### Implementation of dopamine-gated plasticity

Recurrent network modeling approaches typically optimize all parameters **θ** of the network (in this case, **θ** = {**W**_recur_, **W**_KC→MBON_, **W**_readout_, **W**_ext_, **b**}) in order to produce a desired behavior. This approach assumes that, after optimization, connections are fixed to constant values during the execution of the behavior. However, connections between Kenyon cells and output neurons are known to exhibit powerful and rapid dopamine-gated synaptic plasticity. This plasticity is dependent on the relative timing of Kenyon cell and dopamine neuron activation (notably, it does not appear to depend on the postsynaptic output neuron firing rate^26^) and can drive substantial changes in evoked output neuron activity even after brief KC-DAN pairings^28^. We therefore augmented our networks with a model of this plasticity by assuming that each element of **W**_KC→MBON_ is a dynamic quantity that tracks a variable *w* (with a time constant corresponding to the induction of plasticity; see Methods). These variables, which determine the strength of the connection from the *j*th Kenyon cell to the *i*th output neuron, obey the following update rule:

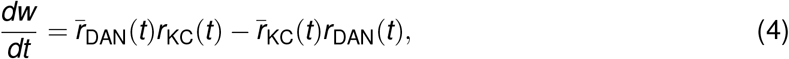

where *r*_KC_ and *r*_DAN_ are the firing rates of the *j*th Kenyon cell and the dopamine neuron that innervates the *i*th compartment, and 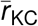 and 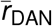 are synaptic eligibility traces constructed by low-pass filtering *r*_KC_ and *r*_DAN_. The time constants of the low-pass filters used to generate the eligibility traces determine the time window within which pairings of Kenyon cell and dopamine neuron activity elicit appreciable changes of *w*. Odors are encoded by sparse activation of random subsets of Kenyon cells, which is accomplished in the model by setting 10% of the elements of **r**_KC_ to 1 and the rest to 0. When Kenyon cell and dopamine neuron firing rates are modeled as pulses separated by a time lag Δ*T*, the dependence of the change in *w* on Δ*T* takes the form of a biphasic timing-dependent function (Fig. 1B,C), consistent with a recent experimental characterization^28^. The seconds-long timescale of this curve is compatible with the use of continuous firing rates rather than discrete spike timing to model KC-to-MBON plasticity, as we have done in Eq. 4.

Importantly, the weight update rule in Eq. 4 is a smooth function of network firing rates, allowing networks with this update rule to be constructed using gradient descent. Specifically, we minimize the loss function Eq. 3 under the assumption that the network follows the dynamics defined by Eq. 1 and Eq. 4. The parameters to be optimized are **θ** = {**W**_recur_, **W**_ext_, **W**_readout_, **b**} (the connections describing the mushroom body output circuitry and the biases), while **W**_KC→MBON_ is treated as a dynamic quantity. Our models are therefore distinguished from typical gradient-descent-optimized recurrent neural network models by the addition a subset of neurons (dopamine neurons) that gate the plasticity of a particular subset of weights (**W**_KC→MBON_) with a timing-dependent rule. This allows us to ask how this plasticity mechanism is employed “online” by our networks to solve behavioral tasks.

We refer to the gradient descent modification of **θ** as the “optimization” phase of constructing our networks. This optimization represents the evolutionary and developmental processes that produce a network capable of efficiently learning new associations^52^. As described above, after this optimization is complete, the output circuitry is fixed but KC-to-MBON weights are subject to synaptic plasticity according to Eq. 4. Our approach therefore separates synaptic weight changes that are the outcome of evolution and development from those due to experience-dependent KC-to-MBON plasticity, which would be conflated if all parameters were optimized with gradient descent (see Supplemental Fig. 1 for a schematic of the optimization and testing phases of our networks). We show that, after optimization, only the latter form of weight update is sufficient to solve the tasks we consider and generalize to related but novel tasks. To begin, we assume that KC-to-MBON weights are set to their baseline values at the beginning of each trial in which new assocations are formed. Later, we will consider the case of continual learning of many associations.

### Models of associative conditioning

We begin by considering models of classical conditioning, which involve the formation of associations between a conditioned stimulus (CS) and unconditioned stimulus (US) such as reward or punishment. A one-dimensional readout of the output neuron population is taken to represent the stimulus valence (Eq. 2), which measures whether the organism prefers (valence > 0) or avoids (valence < 0) the CS. In the model, CS are encoded by the activation of a random ensembles of Kenyon cells. Rewards and punishments are encoded by external inputs to the network that provide input through **W**_ext_ (see Methods).

To construct the model, we optimized the mushroom body output circuitry to produce a target valence in the readout during presentation of CS+ that have been paired with US (first-order conditioning; Fig. 2A,B, top). During presentations of novel CS-US pairings after optimization, this valence is reported for CS+ but not unconditioned stimulus (CS-) presentations. The activities of subsets of model output neurons are suppressed following conditioning, indicating that the network learns to modify its responses for CS+ but not CS-responses (Fig. 2A,B, bottom) This form of classical conditioning requires an appropriate mapping from US pathways to dopamine neurons, but recurrent mushroom body output circuitry is not required; networks without recurrence also produce the target valence (Fig. 2E, top). We therefore considered a more complex set of tasks. Networks were optimized to perform first-order conditioning, to extinguish associations upon repeated presentation of a CS+ without US, and also to perform second-order conditioning.

**Figure 2:**
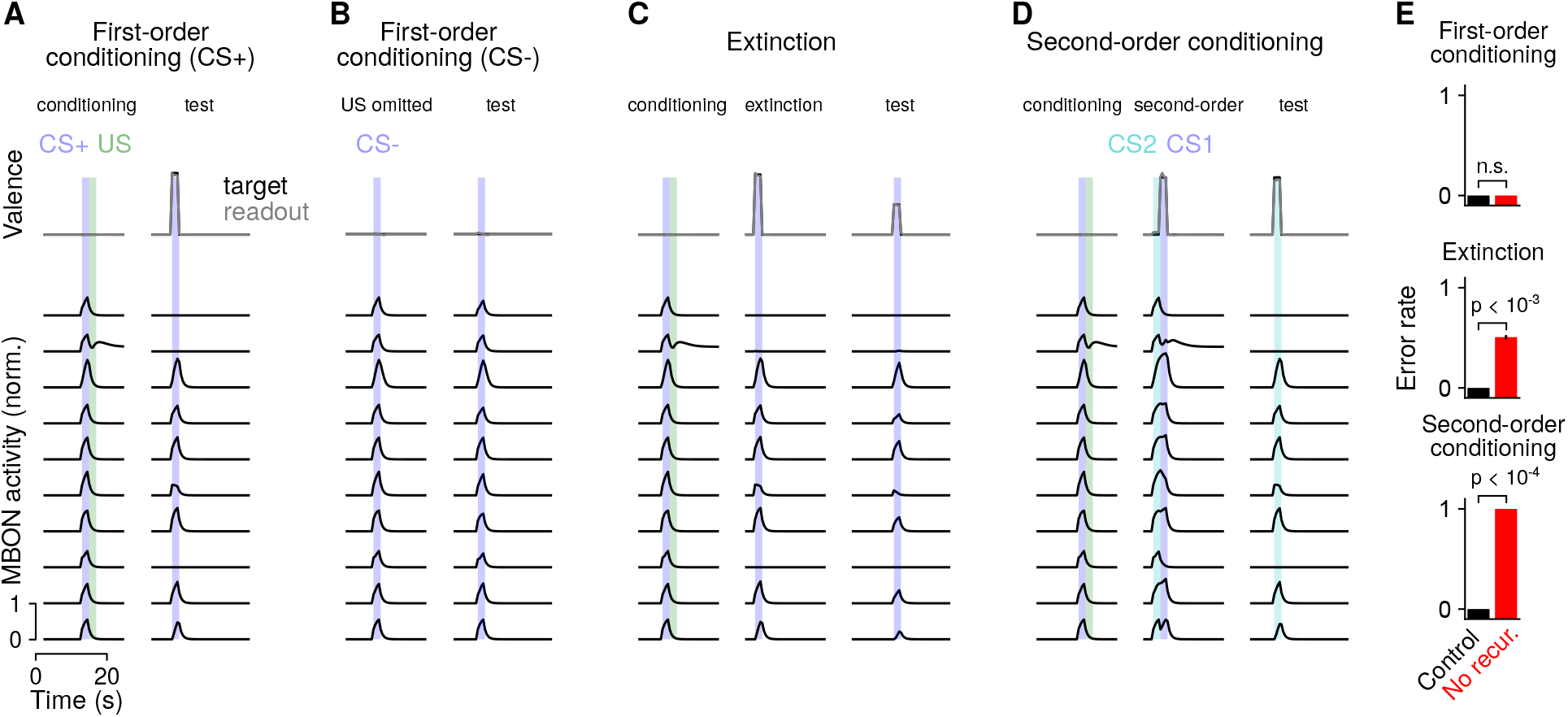
Behavior of network during reward conditioning paradigms. **(A)** Behavior of output neurons (MBONs) during first-order conditioning. During training, a CS+ (blue) is presented, followed by a US (green). Top: The network is optimized so that a readout of the output neuron activity during the second CS+ presentation encodes the valence of the conditioned stimulus (gray curve). Black curve represents the target valence and overlaps with the readout. Bottom: Example responses of output neurons. **(B)** Same as **A**, but for a CS-presentation without US. **(C)** Same as **A**, but for extinction, in which a second presentation of the CS+ without the US partially extinguishes the association. **(D)** Same as **A**, but for second-order conditioning, in which a second stimulus (CS2) is paired with a conditioned stimulus (CS1). **(E)** Error rate averaged across networks in different paradigms. An error is defined as a difference between reported and target valence with magnitude greater than 0.2 during the test period. Networks optimized with recurrent output circuitry (control; black) are compared to networks without recurrence (no recur.; red).

During extinction, the omission of a US following a previously conditioned CS+ reduces the strength of the learned association (Fig. 2C). In second-order conditioning, a CS (CS1) is first paired with a reward or punishment (Fig. 2D, left), and then a second CS (CS2) is paired with CS1 (Fig. 2D, center). Because CS2 now predicts CS1 which in turn predicts reward or punishment, the learned valence of CS1 is transferred to CS2 (Fig. 2D, right). In both extinction and second-order conditioning, a previously learned association must be used to instruct either the modification of an existing association (in the case of extinction) or the formation of a new association (in the case of second-order conditioning). We hypothesized that recurrent output circuitry would be required in these cases. Indeed, non-recurrent mushroom body networks are unable to solve these tasks, while recurrent networks are (Fig. 2E, center, bottom). Thus, for complex relationships between stimuli beyond first-order conditioning, recurrent output circuitry provides a substantial benefit. We also found that these optimized networks generalized to related tasks that they were not optimized for, such as reversal learning (Supplemental Fig. 2), further supporting the conclusion that they implement generalizable learning strategies.

### Comparison to networks without plasticity

Standard recurrent neural networks can maintain stimulus information over time through persistent neural activity, without modification of synaptic weights. This raises the question of whether the dopamine-gated plasticity we implemented is necessary to recall CS-US associations, or if recurrent mushroom body output circuitry alone is sufficient. We therefore compared the networks described above to networks lacking this plasticity. For non-plastic networks, connections from Kenyon cells to output neurons are set to fixed, random values (reflecting the fact that these weights are not specialized to specific odors^53^). These networks evolve similarly to plastic networks except that the dynamics are determined only by Eq. 1 and not by the dopamine-gated plasticity of Eq. 4. Networks are optimized to associate a limited number of CS+ with either a positive or negative valence US, while not responding to CS-. The loss function is the same as Eq. 3.

Non-plastic networks can form CS-US associations (Fig. 3A). Compared to networks with dopamine-gated plasticity (Fig. 2A), output neurons exhibit stronger persistent activity following a CS-US pairing. This activity retains information about the learned association as an “attractor” of neural activity^54^. However, non-plastic networks exhibit a high degree of overgeneralization of learned associations to neutral CS-stimuli (Fig. 3B). This likely reflects a difficulty in constructing a large number of attractors, corresponding to each possible CS-US pairing, that do not overlap with patterns of activity evoked by other CS-stimuli. Consistent with this, as the number of CS+ increases, the difference between the reported valence for CS+ and CS-decreases, reflecting increasing overgeneralization (Fig. 2C). Networks with dopamine-gated plasticity do not suffer from such overgeneralization, as they can store and update the identities of stimuli in plastic weights.

**Figure 3:**
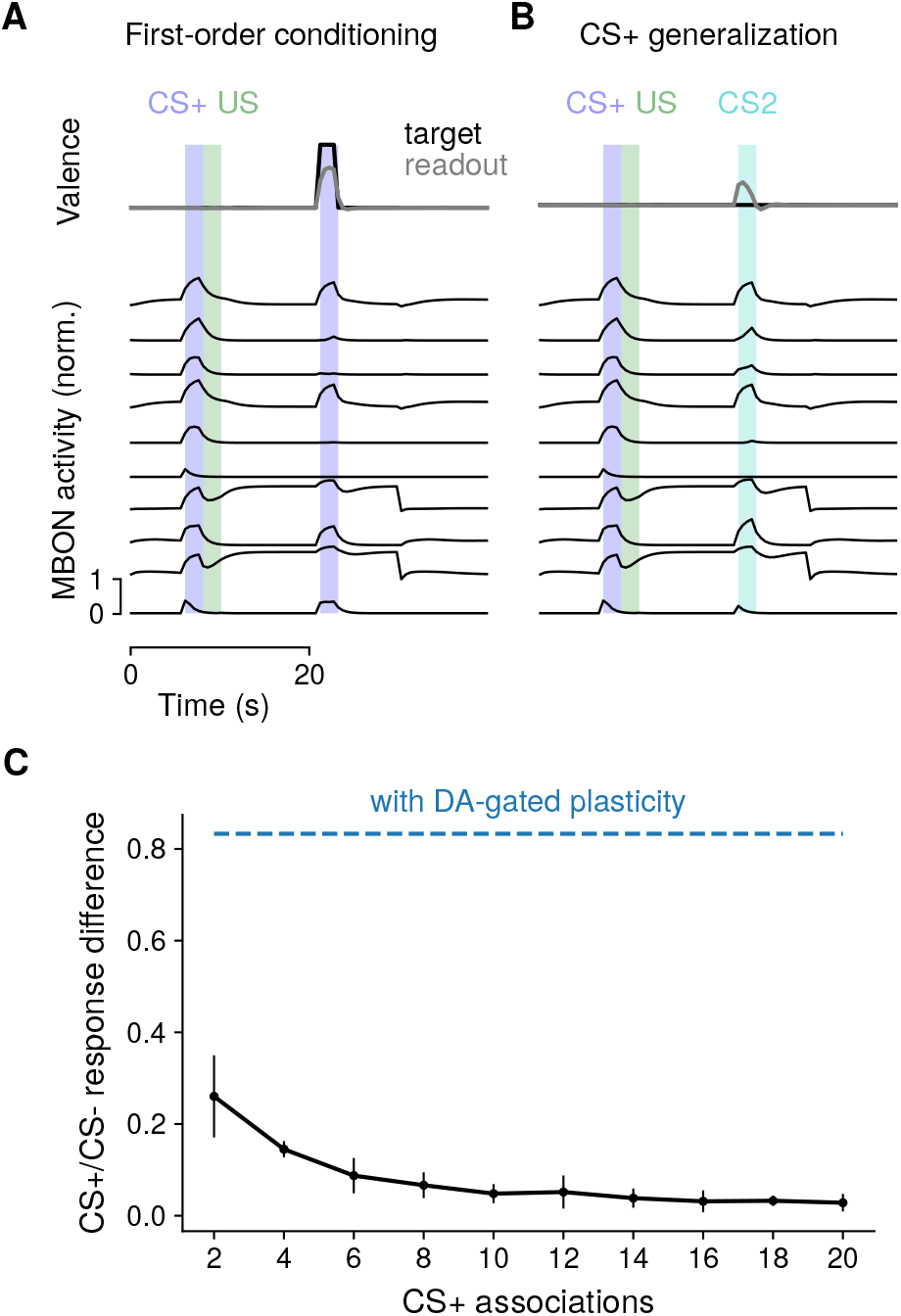
Comparison to networks without dopamine-gated plasticity. **(A)** Behavior during first-order conditioning, similar to Fig. 2A, but for a non-plastic network. Because of the need for non-plastic networks to maintain information using persistent activity, performance degrades with longer delays between training and testing phases. We therefore chose this delay to be shorter than in Fig. 2A. **(B)** Same as **A**, but for a trial in which a CS-US pairing is followed by the presentation of a neutral CS. **(C)** Difference in response (reported valence) for CS+ and CS-as a function of the number of CS+ associations. Each CS+ is associated with either a positive or negative US. A difference of 0 corresponds to overgeneralization of the CS+ valence to neutral CS-. For comparison, the corresponding response difference for networks with dopamine-gated plasticity is shown in blue.

In total, the comparison between plastic and non-plastic networks demonstrates that the addition of dopamine-gated plasticity at KC-to-MBON synapses improves capacity and reduces overgeneralization. Furthermore, plastic networks need not rely solely on persistent activity in order to store associations (compare Fig. 2A and Fig. 3A), likely prolonging the timescale over which information can be stored without being disrupted by ongoing activity.

### Distributed representations across dopamine neurons

We next examined the responses of dopamine neurons to neutral, unconditioned, and conditioned stimuli in the networks we constructed, to examine the “error” signals responsible for learning (Fig. 4A). Dopamine neurons exhibited heterogeneity in their responses. We performed hierarchical clustering to identify groups of dopamine neurons with similar response properties (Fig. 4B, gray; see Methods). This procedure identified two broad groups of dopamine neurons—one that responds to positive-valence US and another that responds to negative-valence US—as well as more subtle features in the population response.

**Figure 4:**
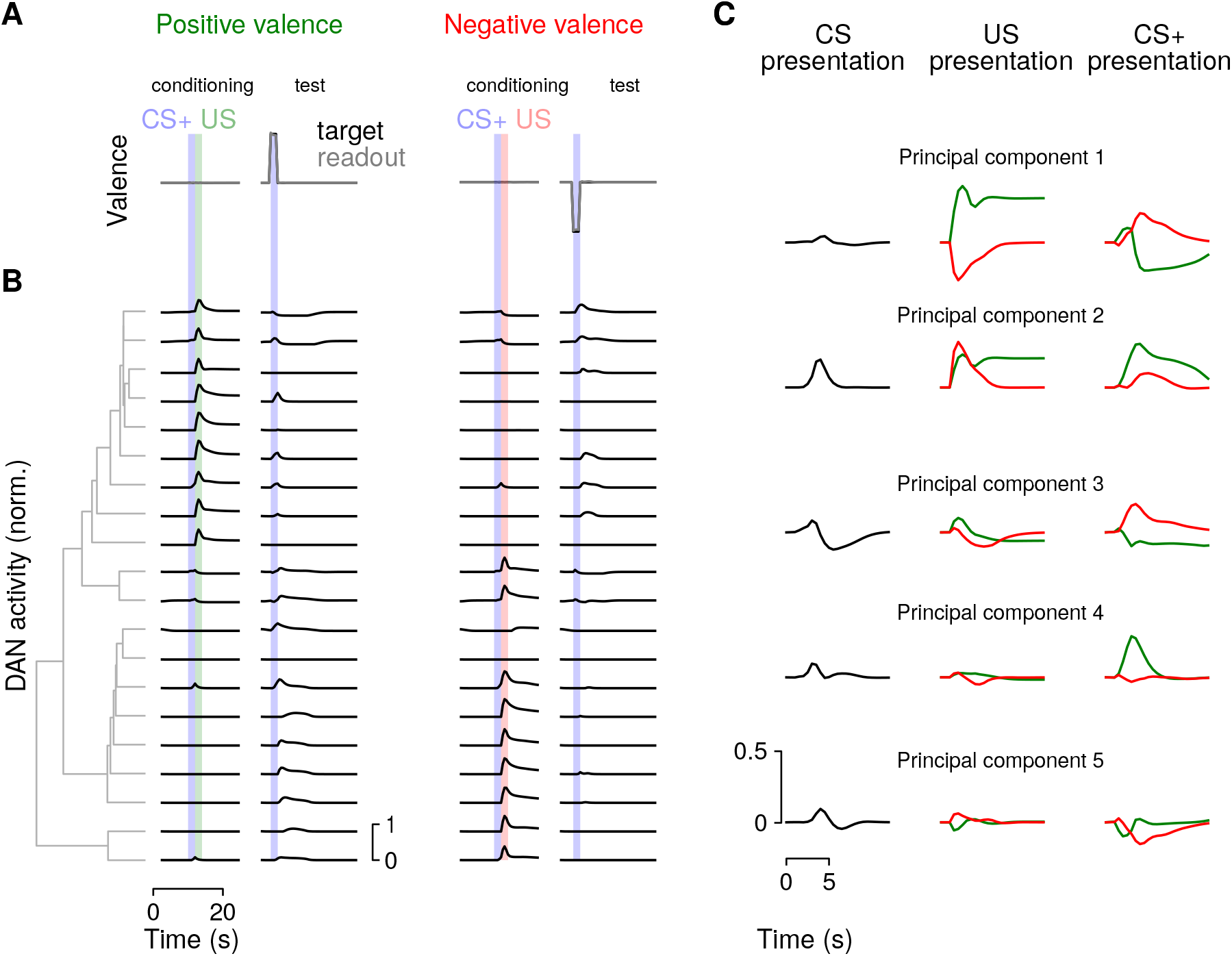
Population analysis of dopamine neuron (DAN) activity. Principal components analysis of dopamine neuron population responses during presentation of neutral CS. **(A)** Responses are shown for CS+ conditioning with a US of positive (left) or negative (valence), followed by a test presentation of the conditioned CS+ without US. **(B)** Responses of model dopamine neurons from a single network. Dopamine neurons are sorted according to hierarchical clustering (illustrated with gray dendrogram) of their responses. **(C)** Principal components analysis of dopamine neuron population activity. Left: Response to a neutral CS. Middle: Response to a positive (green) or negative (red) valence US. Right: Response to a previously conditioned US.

While some dopamine neurons increase their firing only for US, many also respond to reinforced CS. In some cases, this response includes a decrease in firing rate in response to the omission of a predicted US that would otherwise cause an increase in rate, consistent with a reward prediction error. In other cases, neurons respond only with increases in firing rate for US of a particular valence, and for omitted US of the opposite valence, consistent with cross-compartmental interactions supporting the prediction of valence^31^. The presence of both reward prediction error-like responses and valence-specific omission responses suggests that multiple mechanisms are employed by the network to perform tasks such as extinction and second-order conditioning.

The examination of their responses demonstrates that dopamine neurons in our models are diversely tuned to CS and US valence. This tuning implies that KC-to-MBON synapses change in a heterogeneous manner in response to CS and US presentations, but that these changes are sufficient to produce an appropriate behavioral response collectively. Consistent with this idea, principal components analysis of dopamine neuron responses identified modes of activity with interpretable, task-relevant dynamics. The first principal component (Fig. 4C) reflects US valence and predicted CS+ valence, while rapidly changing sign upon US omission, consistent with a reward prediction error. Subsequent principal components include components that respond to CS and US of both valences (principal component 2) or are tuned primarily to a single stimulus, such as a positive valence CS+ (principal component 4).

To further explore how dopamine neuron responses depend on the task being learned, we extended the model to require encoding of novelty and familiarity, inspired by a recent study that showed that the mushroom body is required for learning and expressing an alerting behavior driven by novel CS^29^. We added a second readout that reports CS novelty, in addition to the readout of valence described previously. Networks optimized to report both variables exhibit enhanced CS responses and a large novelty-selective component in the population response identified by principal components analysis (Supplemental Fig. 3), compared to networks that only report valence (Fig. 4B). These results suggest that dopamine neurons collectively respond to any variables relevant to the task for which the output circuitry is optimized, which may include variables distinct from reward prediction. Furthermore, the distributed nature of this representation implies that individual variables may be more readily decoded from populations of dopamine neurons than from single neurons.

### Continual learning of associations

In the previous sections, we modeled the dynamics of networks during individual trials containing a limited number of associations. We next ask whether these networks are capable of continual learning, in which long sequences of associations are formed, with recent associations potentially overwriting older ones. Such learning is often challenging, particularly when synaptic weights have a bounded range, due to the tendency of weights to saturate at their minimum or maximum value after many associations are formed^55^. To combat this, a homeostasic process that prevents such saturation is typically required. We therefore asked if our optimized networks can implement such homeostasis.

In certain compartments of the mushroom body, it has been shown that the activation of dopamine neurons in the absence of Kenyon cell activity leads to potentiation of KC-to-MBON synapses^27^. This provides a mechanism for the erasure of memories formed following synaptic depression.

We hypothesized that this non-specific potentiation could implement a form of homeostasis that prevents widespread synaptic depression after many associations are formed. We therefore augmented our dopamine-gated synaptic plasticity rule (Fig. 1C) with such potentiation (Fig. 5A). The new synaptic plasticity rule is given by:

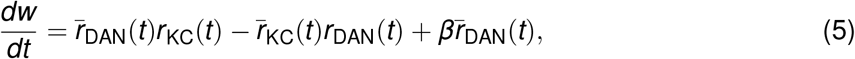

where *β* represents the rate of non-specific potentiation (compare with Eq. 4). We allowed *β* to be optimized by gradient descent individually for each compartment.

**Figure 5:**
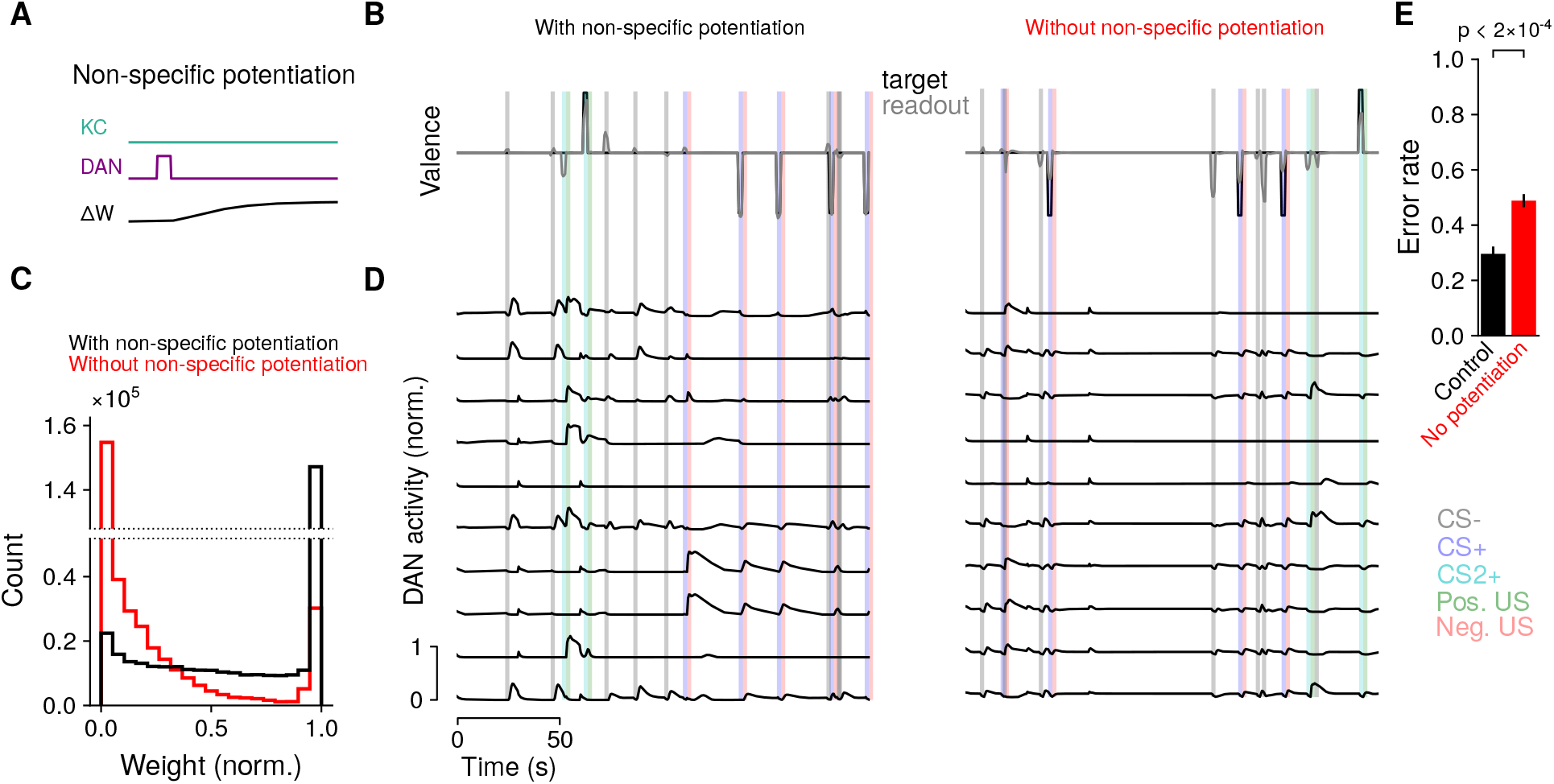
Model behavior for long sequences of associations. **(A)** Illustration of non-specific potentiation following dopamine neuron activity (compare with Fig. 1C). **(B)** Example sequence of positive and negative associations between two odors CS+ and CS2+ and US. Neutral gray odors (CS-) are also presented randomly. **(C)** Histogram of synaptic weights after a long sequence of CS and US presentations for networks with (black) and without (red) non-specific potentiation. Weights are normalized to their maximum value. **(D)** Left: dopamine neuron responses for the sequence of CS and US presentations. Right: same as left, but for a network without non-specific potentiation. Such networks are less likely to report the correct valence for conditioned CS+ and also exhibit a higher rate of false positive responses to CS-. **(E)** Error rate (defined as a difference between reported and target valence with magnitude greater than 0.5 during a CS presentation; we used a higher threshold than Fig. 2 due to the increased difficulty of the continual learning task) for networks with (black) and without (red) non-specific potentiation.

We modeled long sequences of associations in which CS+, CS-, and US are presented randomly (Fig. 5B) and the network is again optimized to produce a target valence (Eq. 3). In optimized networks, the KC-to-MBON weights are initialized at the beginning (*t* = 0) of trial *n* to be equal to those at the end (*t* = *T*) of trial *n* – 1, 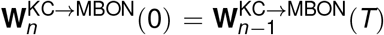, rather than being reset to their baseline values as done previously. We examined the distribution of KC-to-MBON synaptic weights after such sequences of trials. Without non-specific potentiation, most synaptic weights are clustered near 0 (Fig. 5C, red). However, the addition of this potentiation substantially changes the synaptic weight distribution, with many weights remaining potentiated even after thousands of CS and US presentations (Fig. 5C, black).

We also examined performance and dopamine neuron responses in the two types of networks. Without non-specific potentiation, dopamine neuron responses are weaker and the reported valence less accurately tracks the target valence, compared to networks with such potentiation (Fig. 5D,E). In total, we find that our approach can construct models that robustly implement continual learning if provided with homeostatic mechanisms that can maintain a stable distribution of synaptic weights.

### Associating stimuli with changes in internal state

In the previous sections, we focused on networks whose dopamine neurons exhibited transient responses to the presentation of relevant external cues. Recent studies have found that dopamine neurons also exhibit continuous fluctuations that track the state of the fly, even in the absence of overt external reinforcement. These fluctuations are correlated with transitions between, for example, movement and quiescence^33^, or hunger and satiation^56^. Understanding the functional role of this activity is a major challenge for models of dopamine-dependent learning. We hypothesized that such activity could permit the association of stimuli with an arbitrary internal state of the organism. This could allow downstream networks to read out whether a stimulus has previously been experi-enced in conjuction with a particular change in state, which might inform an appropriate behavioral response to that stimulus.

To test this hypothesis, we constructed networks that, in addition to supporting associative conditioning (as in Fig. 2), also transitioned between a set of three discrete internal states, triggered on input pulses that signal the identity of the next state (Fig. 6A). This input represents signals from other brain areas that drive state transitions. We optimized the output circuitry to continuously maintain a state representation, quantified by the ability of a linear readout of dopamine neuron activity to decode the current state (Fig. 6B, top). Specifically, the loss function equaled

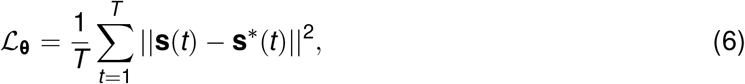

where **s** = Softmax(**W**_readout_**r**_DAN_) is a 3-dimensional vector that represents the decoded probabilities of being in each state and **s**\* is a vector with one nonzero entry corresponding to the actual current state. In this case, **W**_readout_ is a 3 × *N*_DAN_ matrix of weights. Because we were interested in networks that exhibited continuous fluctuations in dopamine neuron activity, we did not impose an additional penalty on dopamine neuron firing rates as in Eq. 3. Optimizing networks with this loss function led to widespread state-dependent activity throughout the network, including among dopamine neurons (Fig. 6B, bottom). This activity coexists with activity evoked by CS or US presentation.

**Figure 6:**
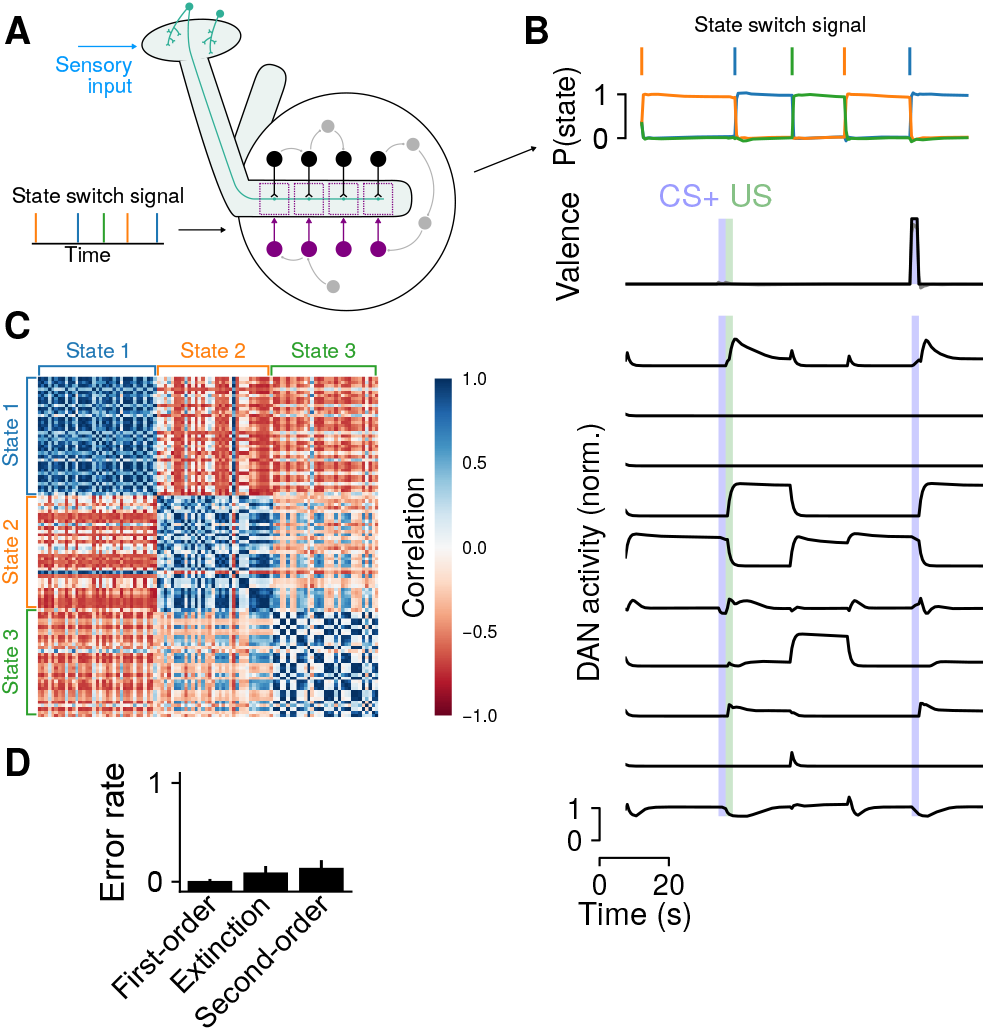
**(A)** Diagram of a network whose activity transitions between a sequence of discrete states in addition to supporting associative conditioning. Brief pulse inputs to the network signal that a switch to a new state should occur. **(B)** Top: A linear readout of dopamine neuron activity can be used to decode the network state. Bottom: dopamine neuron (DAN) activity exhibits state-dependent fluctuations in addition to responding to CS and US. **(C)** Decoding of stimuli that predict state transitions. Heatmap illustrates the correlation between output neuron population responses to the presentation of different stimuli that had previously been presented prior to a state transition. Stimuli are ordered based on the state transitions that follow their first presentation. Blue blocks indicate that stimuli that predict the same state transition evoke similar output neuron activity. **(D)** Performance of networks on conditioning tasks.

We next examined output neuron responses to the presentation of stimuli that had previously preceded a transition to some state. If a transition to a given state reliably evokes a particular pattern of dopamine neuron activity, then KC-to-MBON synapses that are activated by any stimulus preceding such a transition will experience a similar pattern of depression or potentiation. We assessed this response similarity by computing the Pearson’s correlation coefficient 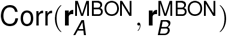, where 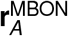 is the average output neuron activity during the presentation of stimulus *A*. Consistent with this prediction, the pattern of output neuron responses evoked by a stimulus that predicts a transition to state *S*_1_ is more similar to the corresponding responses to other stimuli that predict the same state than any other state *S*_2_ (Fig. 6C). The representations of state-transition-predictive stimuli are thus “imprinted” with the identity of the predicted state. While these modifications could potentially interfere with the ability of the system to support associative conditioning, we found that these networks still exhibited high performance on the tasks we previously considered (Fig. 6D). Thus, state-dependent activity and activity required for conditioning are multiplexed in the network. The presence of state-dependent fluctuations could allow circuits downstream of the mushroom body to consistently produce a desired behavior that depends on the internal state, instead of or in addition to the external reinforcement, that is predicted by a stimulus. Our model thus provides a hypothesis for the functional role of state-dependent dopamine neuron activity.

### Mixed encoding of reward and movement in models of navigation

We also examined models of dynamic, goal directed behaviors. An important function of olfactory associations in *Drosophila* is to enable navigation to the sources of reward-predicting odor cues, such as food odors^57^. We therefore optimized networks to control the forward velocity *u*(*t*) and angular velocity *ω*(*t*) of a simulated agent in a two-dimensional environment. We assumed that these movement variables are not decoded directly from output neurons but from other feedback neurons in the mushroom body output circuitry which may represent locomotion-related downstream regions (see Methods). The angular velocity determines the change in the agent’s heading 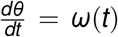, which, along with the forward velocity, determines the change in its location 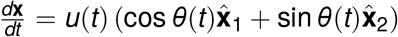 (Fig. 7A). The environment contains multiple odor sources that produce odor plumes that the the agent encounters as it moves. The agent is first presented with a CS+/reward pairing and then is placed in the two-dimensional environment and must navigate to the rewarded odor (Fig. 7A, top). We assumed that the mushroom body output circuitry supports this behavior by integrating odor concentration input from Kenyon cells and information from other brain areas about wind direction relative to the agent’s orientation^58^ (Fig. 7A, bottom; see Methods for a description of how wind input is encoded). Because **x**(*t*) is a differentiable function of network parameters, we can use as a loss function the Euclidean distance between the agent’s location and the rewarded odor source at the end of this navigation period:

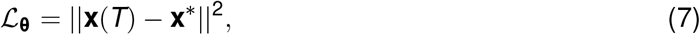

where **x**\* is the location of the rewarded odor source and *T* is the time at which the navigation period ends. Successfully executing this behavior requires storing the identity of the rewarded odor, identifying the upwind direction for that odor, moving toward the odor source using concentration information, and ignoring neutral odors.

**Figure 7:**
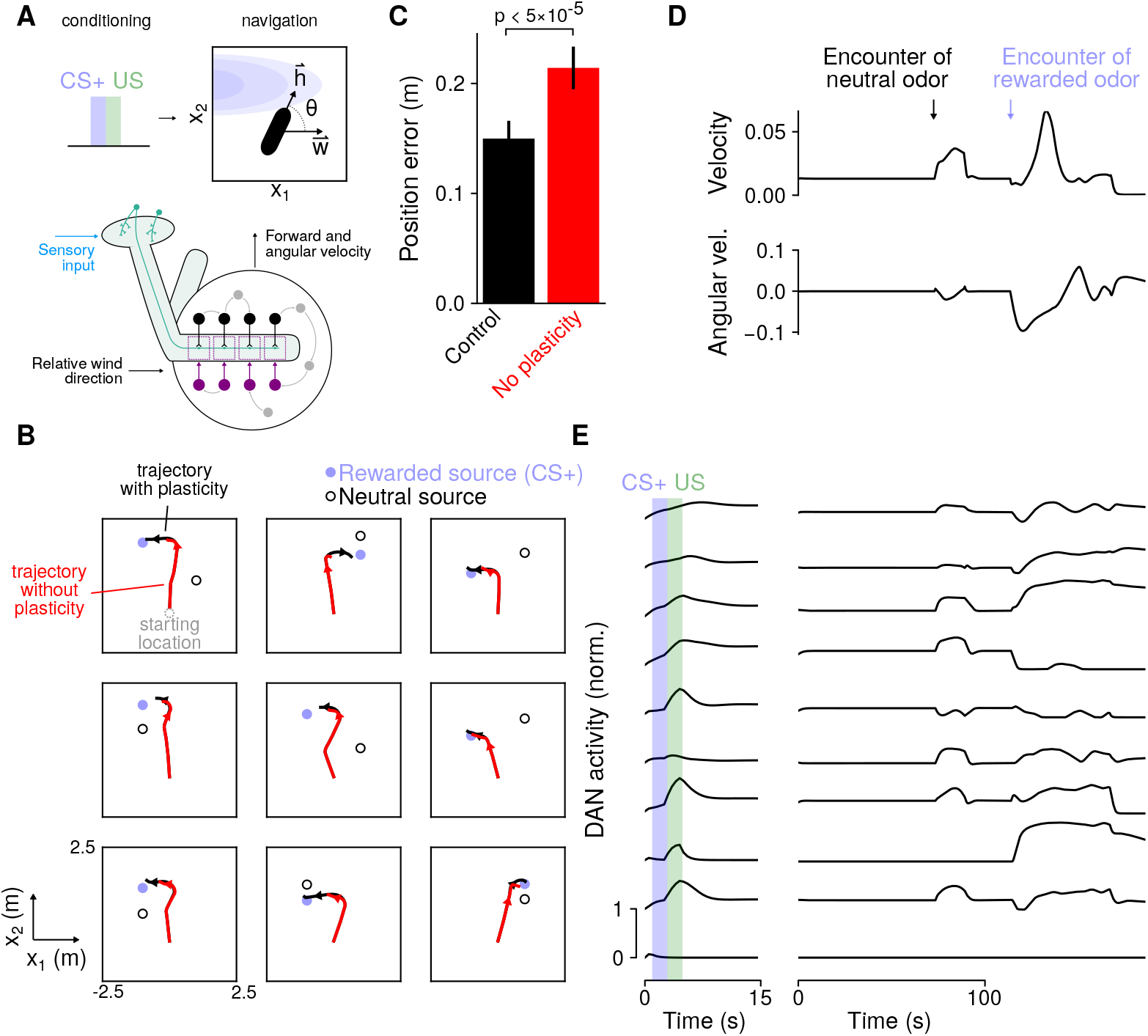
**(A)** Top: Schematic of navigation task. After conditioning, the simulated organism uses odor concentration input (blue) and information about wind direction **w** relative to its heading **h**. Bottom: Diagram of a network that uses these signals to compute forward and angular velocity signals for navigation. Velocity signals are read out from other neurons in the mushroom body output circuitry (gray), rather than output neurons. **(B)** Position of the simulated organism as a function of time during navigation. Black: Simulation with intact dopamine-gated plasticity during navigation; Red: Simulation with plasticity blocked. Arrowheads indicate direction of movement. In the top left plot, the starting location (gray circle) is indicated. **(C)** Position error (mean-squared distance from rewarded odor source at the end of navigation) for control networks and networks without dopamine-gated plasticity. **(D)** Forward (top) and angular (bottom) velocity as a function of time during one example navigation trial. **(E)** Left: Dopamine neuron activity during CS and US presentation in the conditioning phase of a trial. Right: Dopamine neuron activity during the navigation phase of the trial (same trial as in **D**).

The agent can successfully navigate to the rewarded odor source (Fig. 7B), and successful navigation requires plasticity during conditioning that encodes the CS+/US pairing (Supplemental Fig. 4). We wondered whether dopamine-gated plasticity might also be operative during navigation, based on recent findings that recorded ongoing dopamine neuron fluctuations correlated with movement^33^. We asked whether such plasticity during navigation is important for the behavior of the model by examining the performance of networks in which this plasticity is blocked after the networks are optimized. Blocking plasticity during navigation impairs performance, suggesting that it contributes to the computation being performed by the mushroom body output circuitry (Fig. 7C). In particular, networks lacking plasticity often exhibit decreased forward velocity after entering a plume corresponding to a rewarded odor (Fig. 7B), suggesting that ongoing plasticity may reinforce salient odors as they are encountered and promote odor-seeking, consistent with a recent report^59^.

We also examined the relationship between dopamine neuron activity and movement variables during navigation. The agent exhibits increased forward velocity and turning upon encountering an odor, with greater increases for rewarded than for neutral odors (Fig. 7D). Model dopamine neurons exhibit activity during navigation that correlates with movement (Fig. 7E; Supplemental Fig. 5). Many of the same dopamine neurons also exhibit reward-related activity, demonstrating that they multiplex reward and movement-related signals, rather than these classes of dopamine neurons forming disjoint subsets. Thus, our model accounts for dopamine neuron tuning to these two types of signals, a feature present in recordings that traditional modeling approaches do not capture^33^.

## Discussion

We have developed models of the mushroom body that use a biologically plausible form of dopamine-gated synaptic plasticity to solve a variety of learning tasks. By optimizing the mushroom body output circuitry for task performance, these models generate patterns of dopamine neuron activity sufficient to produce the desired behaviors. Model dopamine neuron responses are distributed, tuned to multiple task-relevant variables, and exhibit rich temporal fluctuations. This diversity is a result of optimizing our models only for task performance rather than assuming that dopamine neurons uniformly represent a particular quantity of interest, such as a global reward prediction error signal^3^. Our results predict that individual dopamine neurons may exhibit diverse tuning while producing coherent activity at the population level. They also provide the first unified modeling framework that can account for valence and reward prediction (Fig. 4), novelty (Supplemental Fig. 3), and movement-related (Fig. 7) dopamine neuron responses that have been recorded in experiments.

### Relationship to other modeling approaches

To construct our mushroom body models, we took advantage of recent advances in recurrent neural network optimization to augment standard network architectures with dopamine-gated plasticity. Our approach can be viewed as a form of “meta-learning”^45^, or “learning to learn,” in which a network learns through gradient descent to use a differentiable form of synaptic plasticity (Eq. 4) to solve a set of tasks. As we have shown, this meta-learning approach allows us to construct networks that exhibit continual learning and can form associations based on single CS-US pairings (Fig. 5). Recent studies have modeled networks with other forms of differentiable plasticity, including Hebbian plasticity,^60–62^ but have not studied gated plasticity of the form of Eq. 4. Another recent study examined networks with a global neuromodulatory signal rather than the heterogeneous signals we focus on^63^.

Another recent study used a meta-learning approach to model dopamine activity and activity in the prefrontal cortex of mammals^64^. Unlike our study, in which the “slow” optimization is taken to represent evolutionary and developmental processes that determine the mushroom body output circuitry, in this study the slow component of learning involved dopamine-dependent optimization of recurrent connections in prefrontal cortex. This process relied on gradient descent in a recurrent network of long short-term memory (LSTM) units, leaving open the biological implementation of such a learning process. Like in actor-critic models of the basal ganglia^65^, dopamine was modeled as a global reward prediction error signal.

In our case, detailed knowledge of the site and functional form of plasticity^28^ allowed us to build models that solved multiple tasks simultaneously using only a biologically plausible synaptic plasticity rule. This constraint allows us to predict patterns of dopamine neuron activity that are sufficient for solving these tasks (Fig. 4). Similar approaches may be effective for modeling other brain areas in which the neurons responsible for conveying “error” signals can be identified, such as the cerebellum or basal ganglia^2,66^.

### Heterogeneity of dopamine signaling across species

Dopamine is responsible for a variety of functions in arthropods, including associative memory in honeybees^6^, central pattern generation in the stomatogastric ganglion of lobsters^7^, escape behaviors^8^ and salivation^9^ in the cockroach, and flight production in moths^10^. While dopamine similarly plays many roles in *Drosophila,* including the regulation of locomotion, arousal, sleep, mating^11^, until recently most studies of *Drosophila* mushroom body dopamine neurons have focused on their roles in appetitive and aversive memory formation^12,13,16,18,20–22^. In mammals, while numerous studies have similarly focused on reward prediction error encoding in midbrain dopaminergic neurons^2^, recent reports have also described heterogeneity in dopamine signals reminiscent of the heterogeneity across dopamine neurons in the mushroom body^5,43^. These include reports detailing distinct subtypes of dopamine neurons that convey positive or negative valence signals or respond to salient signals of multiple valences^39,67^, novelty responses^34–38,40^, responses to threat^68^, and modulation of dopamine neurons by movement^41,42^. In many cases, these subtypes are defined by their striatal projection targets, suggesting a compartmentalization of function similar to that of the mushroom body^5^. However, the logic of this compartmentalization is not yet clear.

Standard reinforcement learning models of the basal ganglia, such as actor-critic models, assume that dopamine neurons are globally tuned to reward prediction error signals^65^. Proposals have been made to account for heterogeneous dopamine responses, including that different regions produce prediction errors based on access to distinct state information^69^, or that dopamine neurons implement an algorithm for learning the statistics of transitions between states using sensory prediction errors^70^. Our results are compatible with these theories, but different in that our model does not assume a priori that all dopamine neurons encode prediction errors. Instead, prediction error coding by particular modes of population activity emerges in our model as a consequence of optimizing for task performance (Fig. 4).

### Connecting mushroom body architecture and function

The identification of groups of dopamine neurons that respond to positive or negative valence US^16,24,30,71,72^, output neurons whose activity promotes approach or avoidance^26^, and dopamine-gated plasticity of KC-to-MBON synapses^27,28,73^ has led to effective models of first-order appetitive and aversive conditioning in *Drosophila.* A minimal model of such learning requires only two compartments of opposing valence and no recurrence among output neurons or dopamine neurons. The presence of extensive recurrence^33,46,50,74^ and dopamine neurons that are modulated by other variables^29,31–33^ suggests that the mushroom body modulates learning and behavior along multiple axes.

The architecture of our model reflects the connectivity between Kenyon cells and output neurons, compartmentalization among output neurons and dopamine neurons, and recurrence of the mushroom body output circuitry. While the identities of output neurons and dopamine neurons have been mapped anatomically^46,75^, the feedback pathways have not, so the feedback neurons in our model (gray neurons in Fig. 1A) represent any neurons that participate in recurrent loops involving the mushroom body, which may involve paths through other brain areas. As electron-microscopy reconstructions of these pathways become available, effective interactions among compartments in our model may be compared to anatomical connections, and additional constraints may be placed on model connectivity. By modifying its architecture, our model could be used to test the role of other types of interactions, such as recurrence among Kenyon cells, connections between Kenyon cells and dopamine neurons^47,76^, or direct depolarizing or hyperpolarizing effects of dopamine on output neurons^48^. There is evidence that dopamine-gated synaptic plasticity rules (Fig. 1B) are heterogeneous across compartments, which could also be incorporated into future models^26,27^. While we have primarily focused on the formation of associations over short timescales because the detailed parameters of compartment-specific learning rules have not been described, such heterogeneity will likely be particularly important in models of long-term memory^21,77–81^.

It is unlikely that purely anatomical information, even at the level of a synaptic wiring diagram, will be sufficient to infer how the mushroom body functions^82^. We have used anatomical information and parametrized synaptic plasticity rules along with hypotheses about which behaviors the mushroom body supports to build “task-optimized” models, related to approaches that have been applied to sensory systems^83^. The success of these approaches for explaining neural data relies on the availability of complex tasks that challenge and constrain the computations performed by the models. Therefore, experiments that probe the axes of fly behavior that the mushroom body supports, including behaviors that cannot be described within the framework of classical conditioning, will be a crucial complement to connectivity mapping efforts as models of this system are refined.

## Supplemental figures

**Supplemental Figure 1:**
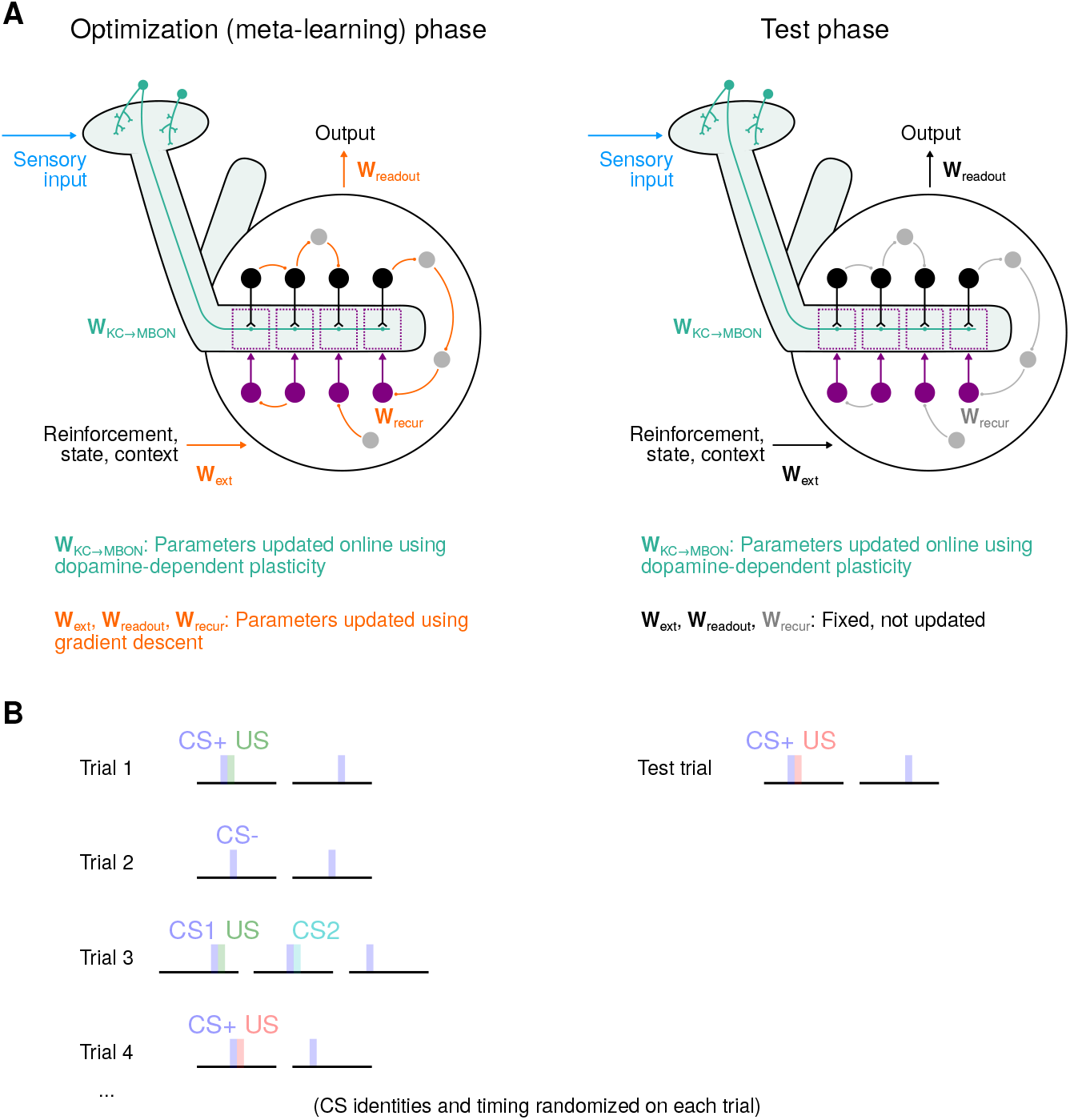
Schematic of meta-learning procedure. **(A)** Two phases of meta-learning and testing. Left: During the optimization phase, connections that form the mushroom body output circuitry are updated with gradient descent (orange). Kenyon cell to output neuron weights evolve “online” (within each trial) according to dopamine-dependent synaptic plasticity. Right: After optimization is complete, the network is tested on a new set of trials. In this phase, connections that form the output circuitry are fixed. **(B)** Illustration of trials involving CS/US associations presented during training (left) and testing (right). Each trial involves new CS/US identities and timing.

**Supplemental Figure 2:**
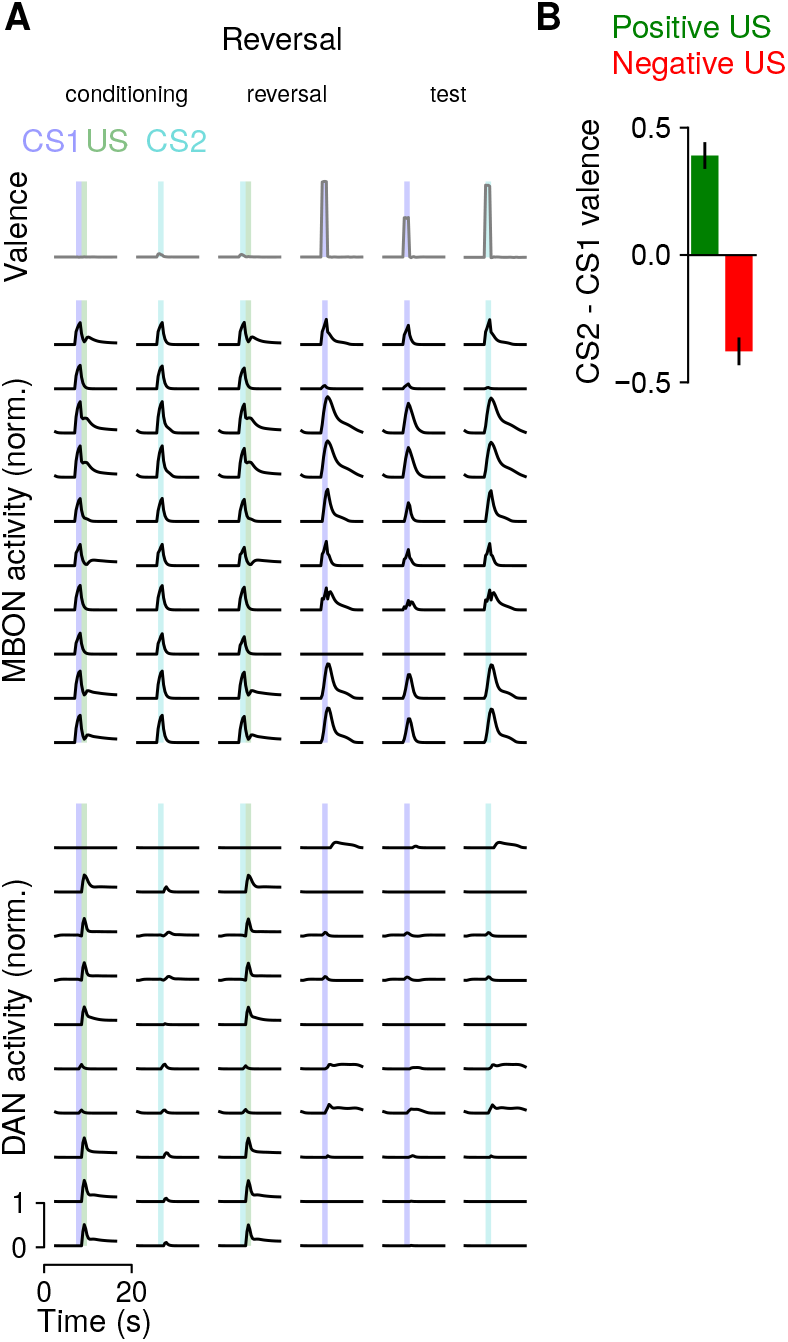
Behavior of networks optimized to perform classical conditioning on a reversal learning task. **(A)** Top: Schematic of reversal learning task. In the first phase, CS1 but not CS2 is paired with US, while during reveral the contingencies are reversed. Preference between CS1 and CS2 is compared in the test phase. Bottom: Example MBON and DAN activity during reversal learning. **(B)** The average difference in reported valence for CS2 vs. CS1. Positive or negative values for positive or negative-valence US, respectively, indicate successful reversal learning. Bars indicate standard deviation across model networks.

**Supplemental Figure 3:**
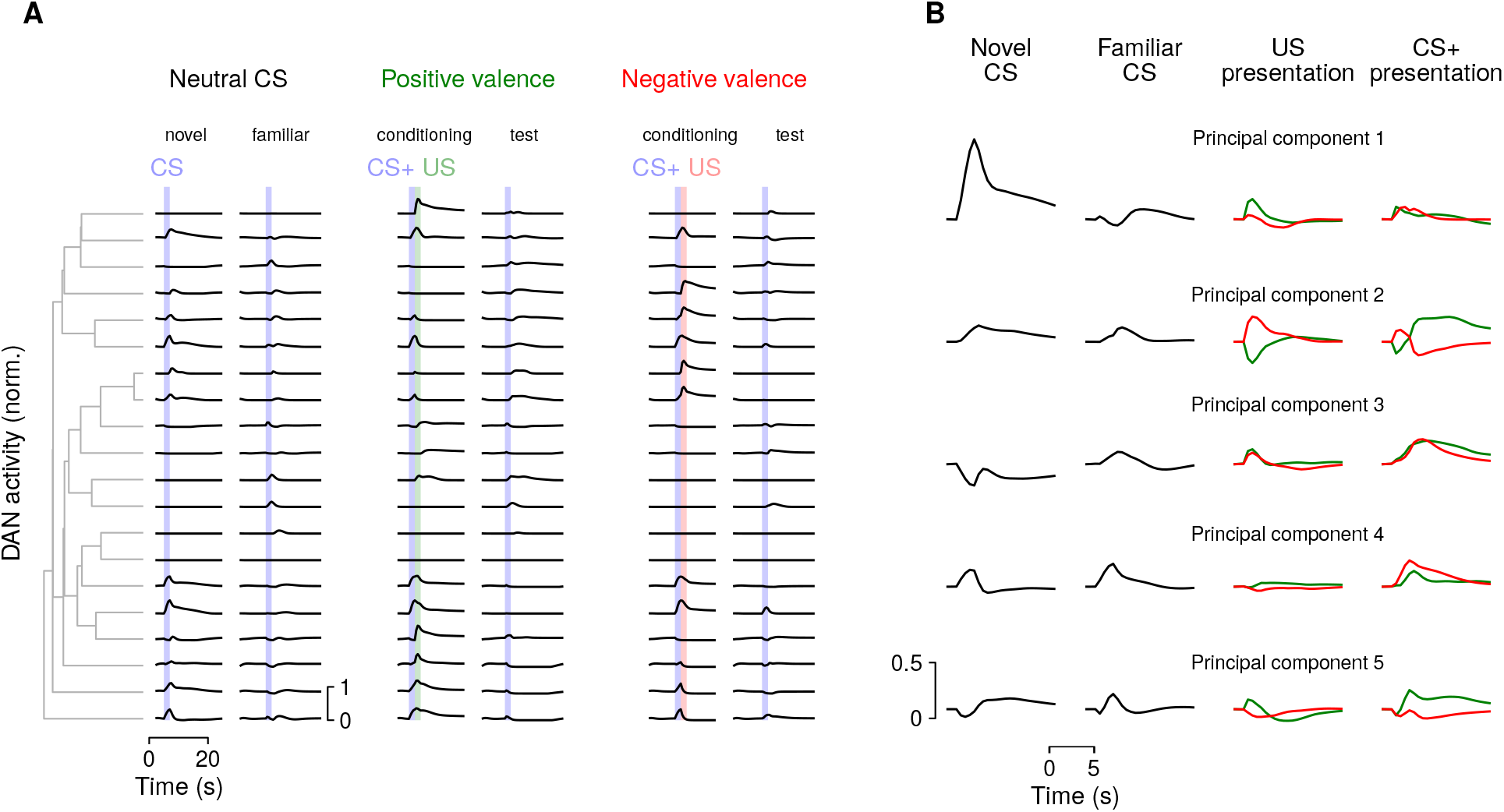
Similar to Fig. 4, but for a network with two readouts that encode both valence and novelty. The novelty readout is active for the first presentation of a given CS and zero otherwise. **(A)** The addition of novelty as a readout dimension introduces dopamine neuron responses that are selective for novel CS. Compare with Fig. 4B. **(B)** The first principal component (PC1) for the network in **A** is selective for CS novelty. Compare with Fig. 4C.

**Supplemental Figure 4:**
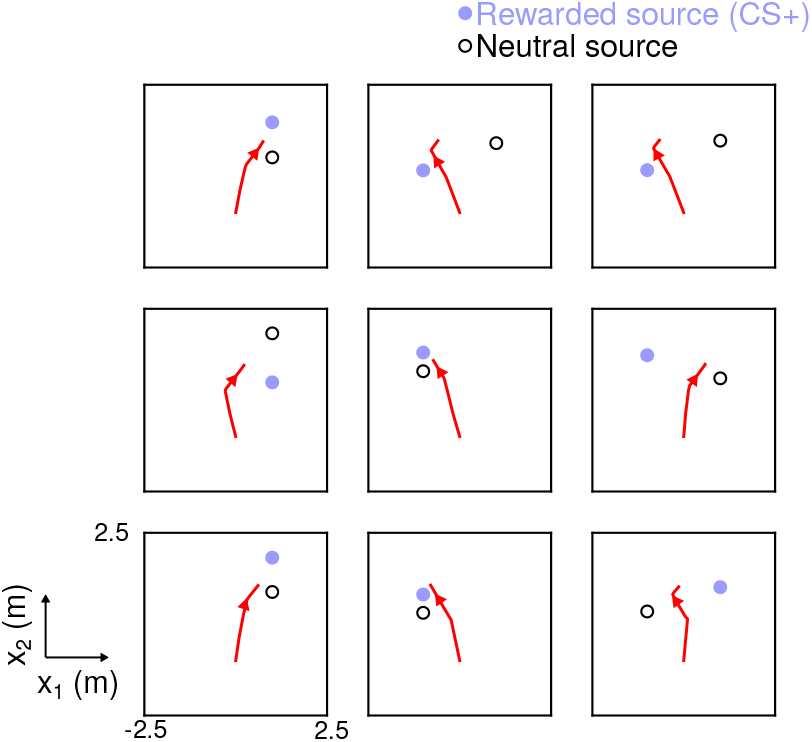
Similar to Fig. 7B, but for a network lacking KC-to-MBON synaptic plasticity during both conditioning and navigation. The model organism is unable to identify the rewarded odor and navigate toward it. Trajectories tend toward points located between the two odor sources.

**Supplemental Figure 5:**
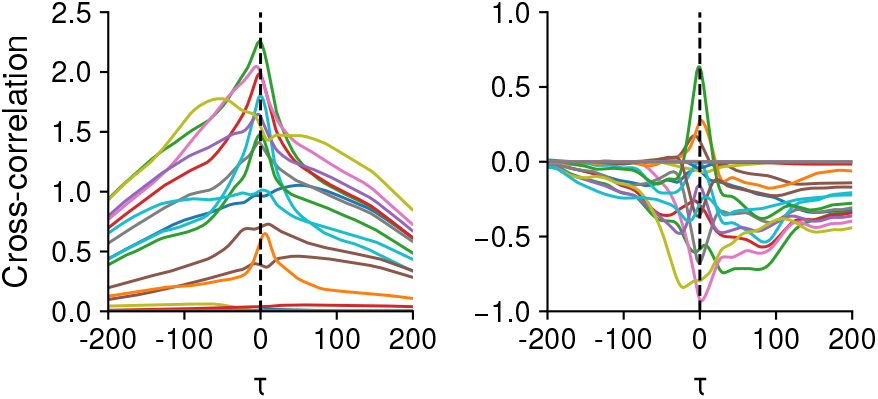
Example cross-correlation functions between dopamine neuron activity and velocity during navigation. Left: Expectation of *d*(*t*)*u*(*t* + *τ*), where *d*(*t*) is dopamine neuron activity and *u*(*t*) is forward velocity. Each color represents a different dopamine neuron. Right: Same as left, but for *d*(*t*)*ω*(*τ* + *τ*), where *ω*(*t*) is angular velocity.

## Methods

### Network dynamics

The dynamics of the networks are determined by Eq. 1 and Eq. 4. If neuron *i* is an output neuron, then its external input is given by 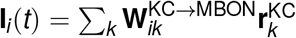, representing input from Kenyon cells. If neuron *i* is a feedback neuron (FBN), then 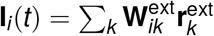, representing reinforcement, context, or state-dependent input from other brain regions. For dopamine neurons, **I**_*i*_(*t*) = 0, as we assume that all input to the dopamine neurons is relayed by feedback neurons..

For KC-to-MBON synapses, each weight is initially set to its maximum value of 0.05 and subsequently updated, with the updates of **W**_KC→MBON_ low-pass filtered with a timescale of *τ_W_* = 5 s to account for the timescale of LTD or LTP. Specifically,

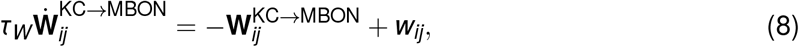

where the weight *w_ij_* for a connection from the *j*th Kenyon cell to the output neuron in compartment *i* is determined by Eq. 4.

KC-to-MBON weights are constrained to lie between 0 and 0.05. For computational efficiency and ease of training, we assume *τ* in Eq. 1 is equal to 1 s and simulate the system with a timestep of Δ*t* = 0.5 s, but our results do not depend strongly on these parameters.

### Optimization

Parameters are optimized using PyTorch with the RMSprop optimizer^84^ (www.pytorch.org) and a learning rate of 0.001. The loss to be minimized is described by Eq. 3, Eq. 6, or Eq. 7 for networks optimized for conditioning tasks, continuous state representations, or navigation respectively. Optimization is performed over a set number of epochs, each of which consists of a batch of *B* = 30 trials. The loss 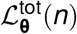 for epoch *n* is the average of the individual losses over each trial in the batch:

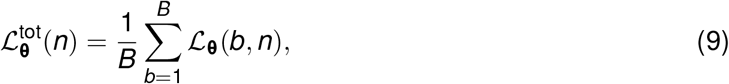

where 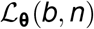 represents the loss for *b*th trial drawn on epoch *n*.

All optimized weights are initialized as zero mean Gaussian variables. To initialize **W**_recur_, weights from a neuron belonging to neuron type *X* (where *X* = MBON, DAN, or FBN) have 0 mean and variance equal to 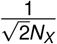, where *N_X_* equals the number of neurons of type *X*. For **W**_readout_, the variance is 1 /*N*_MBON_ while for **W**_ext_, the variance is 1. Bias parameters are initialized at 0.1. At the beginning of each trial, firing rates are reset to an initial state **r**_0_, with **r**_0_ = 0 for output neurons and 0.1 for dopamine neurons or feedback neurons, to permit these neurons to exhibit low levels of baseline activity.

### Conditioning tasks

For conditioning tasks in which the predicted valence of a conditioned stimulus (CS) is reported (such as first- and second-order conditioning and extinction), each CS is encoded by setting 10% of the entries of **r**_KC_ to 1 and the rest to 0. Unconditioned stimuli (US) are encoded by **r**_ext_ which is equal to (1,0)^*T*^ when a positive-valence US is present, (0,1)^*T*^ when a negative-valence US is present, and (0,0)^*T*^ otherwise. CS and US are presented for 2 s. Tasks are split into 30 s intervals (for example conditioning and test intervals; see Fig. 2). Stimulus presentation occurs randomly between 5 s and 15 s within these intervals. Firing rates are reset at the beginning of each interval (e.g. **r**(*t* = 30 s) = **r**_0_), which prevents networks from using persistent activity to maintain associations.

When optimizing networks in Fig. 2, random extinction and second-order conditioning trials were drawn. For half of these trials, CS or US are randomly omitted (and the target valence updated accordingly - e.g., if the US is omitted, the network should not report a nonzero valence upon the second CS presentation; Fig. 2B) in order to prevent the networks from overgeneralizing to unconditioned CS. Optimization progressed for 5000 epochs for networks trained to perform extinction and second-order conditioning. For networks trained only for first-order conditioning, (Fig. 2E, top; Fig. 3), only first-order conditioning trials were drawn, and optimization progressed for 2000 epochs.

Principal components of dopamine neuron activity (Fig. 4) were estimated using 50 randomly chosen trials of extinction and second-order conditioning in previously optimized networks. To order dopamine neurons based on their response similarity (Fig. 4A), hierarchical clustering was performed using the Euclidean distance between the vector of firing rates corresponding to pairs of dopamine neurons during these trials.

For networks also trained to report stimulus novelty (Supplemental Fig. 3), an additional readout dimension *n*(*t*) that is active for the first presentation of a given CS and inactive otherwise is added. The full network readout is then given by

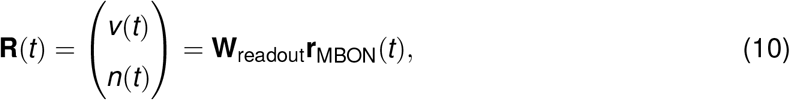

and the loss equals

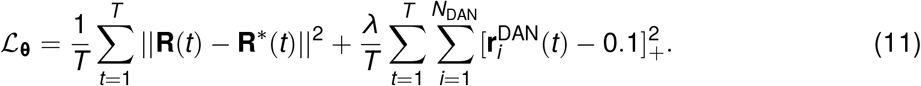

Adding this second readout does not significantly impact the performance of the networks for classical conditioning tasks.

### Networks without dopamine-gated plasticity

For networks without dopamine-gated plasticity, KC-to-MBON synaptic weights are drawn randomly from a uniform distribution between 0 and 0.05 and then fixed. The time of CS+ presentation is chosen uniformly between 5 s and 15 s, and the second CS presentation occurrs uniformly between 20 s and 30 s. Networks are optimized to perform first-order conditioning with positive and negative valence US for a fixed set of CS+ stimuli numbering between 1 and 10 (2 to 20 possible associations). On half of the trials, a random CS is presented instead of the second CS+ presentation (Fig. 3B) and networks are optimized to not respond to this CS.

### Continual learning

To model continual learning (Fig. 5), networks were augmented with non-specific potentiation gated by dopamine neuron activity according to Eq. 5. The potentiation parameter *β* is compartment-specific and updated through gradient descent. Each parameter is initialized at 0.01 and constrained to be positive.

Trials consist of 200 s intervals, during which two CS+ and two CS-odors are presented randomly. For each CS, the number of presentations in this interval is chosen from a Poisson distribution with a mean of 2 presentations. Unlike other networks, for these networks the values of **W**_KC→MBON_ at the end of one trial are used as the initial condition for the next trial. To prevent weights from saturating early in optimization, the weights at the beginning of trial *t* are set equal to:

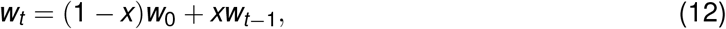

where *w*_0_ = 0.05 corresponds to the initial weight at the beginning of optimization, and *x* increases linearly from 0 to 1 during the first 2500 epochs of optimization. Networks were optimized for a total of 5000 epochs.

### Networks that encode changes in state

For networks that encode changes in state (Fig. 6), we modified our training protocol for networks optimized for conditioning tasks to include an additional three-dimensional readout of output neuron activity optimized to encode the state (at each moment in time, the target is equal to 1 readout dimension and 0 for the others; Eq. 6). The external input **r**_ext_ is three-dimensional and signals state transitions using input pulses of length 2 s. The length of time between pulses Δ*T*_state_ is a random variable distributed according to Δ*T*_state_ ~ 10 s o (1 + Exp(1)). Networks were optimized for 500 epochs.

To test how state-dependent dopamine neuron dynamics affect stimulus encoding, a CS is presented for 2 s, beginning 8 s prior to the second state change of a 300 s trial. Afterward, the same CS is presented for 5 s. This was repeated for 50 CS, and the correlation coefficient between output neuron responses during the second 5 s presentation was calculated (Fig. 6C).

### Models of navigation

To model navigation toward a rewarded odor source (Fig. 7), a CS+/US pairing is presented at *t* = 2 s in a 20 s training interval with a US strength of 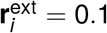. This is followed by a 200 s interval during which the model organism navigates in a two-dimensional environment.

During navigation, two odor sources are present, one CS+ and one neutral CS. The sources are randomly placed at *x*_1_ = ±1 m and *x*_2_ chosen uniformly between 0 m and 2 m, with a minimum spacing of 0.5 m. Associated with each odor source is a wind stream that produces an odor plume that the model organism encounters as it navigates. These are assumed to be parallel to the *x*_1_ axis and oriented so that the odor plume diffuses toward the origin, with a height of 0.5 m and centered on the *x*_2_ position of each odor source. For locations within these plumes and downwind of an odor source, the concentration of the odor is given by:

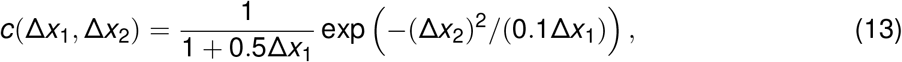

where Δ*x*_1_ and Δ*x*_2_ are the *x* and *y* displacements from the odor source in meters. This equation expresses a Gaussian odor plume with a width that increases and magnitude that decreases with distance from the odor source.

During navigation, when the model organism encounters an odor plume, Kenyon cell activity is assumed to be proportional to the pattern of activity evoked by an odor (as before, a random pattern that activates 10% of Kenyon cells) scaled by *c*(Δ*x*_1_, Δ*x*_2_). The network further receives 4-dimensional wind direction input via **W**_ext_. Each input is given by [**w** · **h**_*i*_]_+_, where **w** is a unit vector representing wind direction and **h**_*i*_ for *i* = 1 … 4 is a unit vector pointing in the anterior, posterior, or lateral directions with respect to the model organism.

The organism is initially placed at the origin and at an angle distributed uniformly on the range 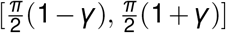, with *γ* increasing linearly from 0 to 0.5 during the optimization. The movement of the organism is given by two readouts of the feedback neurons. The first determines the forward velocity *u*(*t*) = Softplus(**W**_*u*_ · **r**(*t*)+ *b_u_*), and the second determines the angular velocity *ω*(*t*) = **W**_*ω*_ · **r**(*t*) + *b_ω_*. The weights and bias parameters of these readouts are included in the parameter vector **θ** that is optimized using gradient descent. For each trial, the loss is determined by the Euclidean distance of the model organism from the rewarded odor source at the end of the navigation interval (Eq. 7). Networks were optimized for 500 epochs.

### Code availability

Code implementing the models will be made available upon publication.

## Acknowledgments

We wish to thank L. F. Abbott, Y. Aso, R. Axel, V. Ruta, M. Zlatic, and A. Cardona for insightful discussions and comments on the manuscript. We are particularly grateful to L. F. Abbott for discussions during the development of this study. Research was supported by a Columbia University Class of 1939 Summer Research Fellowship (L. J.), the Columbia Science Research Fellows Program (L. J.), the Burroughs-Wellcome Foundation (A. L.-K.), the Simons Collaboration on the Global Brain (A. L.-K.), the Gatsby Charitable Foundation (L. J. and A. L.-K.), and NSF NeuroNex Award DBI-1707398 (L. J. and A. L.-K.).

